# MSImpute: Imputation of label-free mass spectrometry peptides by low-rank approximation

**DOI:** 10.1101/2020.08.12.248963

**Authors:** Soroor Hediyeh-zadeh, Andrew I. Webb, Melissa J. Davis

**Affiliations:** Bioinformatics Division, Walter and Eliza Hall Institute of Medical Research, VIC, Australia; Systems Biology and Personalized Medicine Division, Walter and Eliza Hall Institute of Medical Research, VIC, Australia; Department of Medical Biology, University of Melbourne, VIC, Australia; Department of Clinical Pathology, Faculty of Medicine, Dentistry and Health Sciences, University of Melbourne, VIC, Australia

## Abstract

Recent developments in mass spectrometry (MS) instruments and data acquisition modes have aided multiplexed, fast, reproducible and quantitative analysis of proteome profiles, yet missing values remain a formidable challenge for proteomics data analysis. The stochastic nature of sampling in Data Dependent Acquisition (DDA), suboptimal preprocessing of Data Independent Acquisition (DIA) runs and dynamic range limitation of MS instruments impedes the reproducibility and accuracy of peptide quantification and can introduce systematic patterns of missingness that impact downstream analyses. Thus, imputation of missing values becomes an important element of data analysis. We introduce `msImpute`, an imputation method based on low-rank approximation, and compare it to six alternative imputation methods using public DDA and DIA datasets. We evaluate the performance of methods by determining the error of imputed values and accuracy of detection of differential expression. We also measure the post-imputation preservation of structures in the data at different levels of granularity. We develop a visual diagnostic to determine the nature of missingness in datasets based on peptides with high biological dropout rate and introduce a method to identify such peptides. Our findings demonstrate that `msImpute` performs well when data are missing at random and highlights the importance of prior knowledge about nature of missing values in a dataset when selecting an imputation technique.

## 1 Introduction

Measurement of protein abundance in samples provides vital insight into the molecular processes that underpin biological or clinical phenotypes of interest. Label-free liquid chromatography mass spectrometry (LC-MS) is commonly used for comprehensive characterization of protein species in a sample (*1*), yet a substantial fraction of identified peptide measurements is missing from proteomics datasets (*2*). In proteomic biomarker discovery studies, where no prior information is available on the potential protein species that exist in a sample, data is acquired by the spectrum centric mode data dependent acquisition (DDA). The stochastic sampling of peptide features for fragmentation during DDA combined with low signal to noise ratio at lower levels near to limit of detection of MS instruments and imperfect feature detection result in high levels of missing values, even within technical replicates (*3*). Alternatively, the peptide centric data independent acquisition (DIA) mode of operation requires prior knowledge about the fragment ion spectra of targeted peptides and has substantially enhanced the reproducibility of proteome quantification (*4*). However missing values still remain a problem for the field, and can impact results from the downstream statistical analysis of proteomics data (*5*).

Missing values can be categorized into three categories: Missing Completely At Random (MCAR), Missing At Random (MAR) and Missing Not At Random (MNAR) (*6*). MCAR missing values in proteomics data can originate from random errors or stochastic fluctuations during experimental process. A number of different factors are reported to impact the accuracy and reproducibility including, sample preparation, sample processing, peptide separation, changes in sample complexity, matrix effects and ion suppression, detector saturation and other technical factors (*7–10*). MAR is defined as the possibility of a variable being missing is dependent on other observed variables (*6, 11*). MAR data in proteomics are produced during data preprocessing, for example, by inaccurate peak detection and deconvolution of co-eluting compounds. Missingness of values under the limits of quantification (LOQ) (i.e. left-censored missing) are considered as MNAR (*2*).

The imputation methods in proteomics are broadly categorized as single value approaches, local similarity and global similarity approaches (*12*). Imputation of left-censored MNAR missing values is typically performed by replacing the missing value with a small value or with zero, although more sophisticated methods such as quantile regression imputation have also been applied to left-censored data. MAR/MCAR are generally difficult to distinguish and can be imputed by local methods based on observed values in the neighborhood, or global similarity approaches such as maximum likelihood estimation (*13*). We propose a global imputation method that completes missing values by estimating a low-rank representation of the data and evaluate this method against common imputation approaches in simulated and real MNAR and MAR/MCAR data. We also provide a diagnostic for MAR/MNAR in proteomics data based on peptides with higher than expected dropout rate, and demonstrate how this diagnostic can guide the selection of imputation method.

## 2 Experimental Procedures

We propose a method for imputation of missing peptide intensity measurements in label-free proteomics experiments. Assuming that peptide intensities profiled in several samples in an experiment are represented as a matrix, where rows are peptides and columns correspond to samples, the proposed method re-constructs the partially observed matrix of peptide intensities by estimating the underlying patterns in abundance data; a low-rank approximation to the incomplete peptide intensity matrix. We benchmark the proposed method in several public datasets, evaluate the performance of the method along with other popular imputation procedures in simulated data, and provide parallel evidence that support our findings in empirical datasets. Additionally, we propose a model that serves as a visual diagnosis to deduce the MAR/NMAR nature of missing values in datasets.

### 2.1 Experimental Design and Statistical Rationale

#### 2.1.1 Benchmark datasets

A number of public DDA and DIA benchmark datasets with Universal Proteomics Standards (UPS1/2) or known spiked-in proteins were identified from the previous literature. An overview of the datasets used in this study is given in Table 1.

**Table 1.** Datasets used in this study, listed in the order of appearance in figures.

| Dataset name | Description | Type | No. of replicates | Reference |
| --- | --- | --- | --- | --- |
| PXD011691 | Samples from 10 different mouse cerebelli spiked with the UPS2 protein standard in five different concentrations. | DIA, TMT 10-plex | 10 | Muntel et al.<sup>17</sup> |
| PXD002370 | Three different concentrations of UPS1 (Sigma) spiked in yeast extracts. | DDA | 3 | Giai Gianetto et al.<sup>15</sup> |
| PASS00589 | Three master mixes consisting of twelve non-human proteins spiked into a constant background (HEK-293) at various concentrations. | DDA, DIA | 4 | Bruderer et al.<sup>16</sup> |
| PXD000279 | Dynamic range benchmark dataset consisting of UPS1/UPS2 standards (Sigma) spiked into E. coli lysates. | DDA | 4 | Cox et al.<sup>14</sup> |

**PXD000279** consists of UPS1/UPS2 standards spiked into E.coli lysates. Four replicates are available for each UPS Proteomics Standards. This is a DDA dataset (*14*).

**PXD002370** is a DDA dataset, where three concentrations of UPS1 (25 fmol, 10 fmol and 5 fmol) were spiked in yeast extract in triplicates. PXD002370-Dataset 1 is a comparison between 10 fmol and 5 fmol, and PXD002370-Dataset 2 compares 25 fmol to 10 fmol (*15*).

**PASS00589** consists of a DDA and a DIA dataset, where a set of 12 proteins are spiked at different concentrations. Each dataset comes in four biological and three technical replicates (*16*).

**PXD011691** The sample set in PXD011691 consisted of ten samples derived from 10 different mouse cerebelli spiked with the UPS2 protein standard in five different concentrations (*17*). It contains DIA and TMT datasets acquired at two facilities. We used the DIA dataset from FLI facility in differential expression analysis. We used the DIA and TMT datasets from BGS center in the empirical evaluation of RMSE and local/global structures, and the TMT data for generation of missing values, as it has smallest proportion of missing values compared to data from FLI facility. We only retained complete observations in the TMT data from BGS facility prior to amputation (a process where we mask a real, measured value to treat it as if it were missing, allowing us to impute a value in a situation where we know the real, observed value).

The following pre-processing steps were applied to peptide-level intensities: contaminant peptides or those assigned to the reverse sequence of a known peptide or decoy peptides were removed. Signal intensity values were collapsed at the peptide sequence level by adding signal intensities from different charge states together. Peptides were retained only if they were measured (not NA) in more than four samples. Peptides are typically filtered if they are not detected in at least k samples, where k is the minimum number of biological replicates in an experimental group. However, low-rank approximation requires at least 4 observed measurements per peptide. Peptide intensities were then log_2_-transformed and quantile normalized. If a quantification workflow reports missing values as zeros, the values were set back to NAs. Datasets were imputed at the peptide level.

#### 2.1.2 Missing value generation - data simulation

We utilized the `ampute` function (*18–20*) in *mice* R package (*21*) to simulate MAR and MNAR missingness patterns in TMT dataset from (*17*) with 10 samples. The function expects a complete dataset, and a list of peptides (a combination of which is called a pattern) selected for amputation (masking), the fraction of samples affected by the pattern (referred to as pattern frequency herein), a mechanism for missing values, and proportion of missing values. Peptides were selected for amputation based on a mean-dependent, weighted sampling strategy, such that peptides at low abundance receive higher weight for being selected for amputation, whereas less weight is assigned to peptides at high abundance. This mean-dropout trend is previously reported by (*12*), and can be modeled by the following sampling strategy. Let *µ*_i_ denote the standardized average log_2_ intensity for peptide *i* across all peptides. Each peptide is selected according to a weight defined by

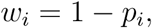

where *p*_*i*_ is computed from a logistic function

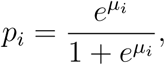

For MAR generation, we selected 30% of the peptides according to the sampling procedure described above. The selected peptides were partitioned into three patterns. Each pattern defines a combination of peptides selected for amputation. The first pattern was set to be missing in 60% of samples, and the two remaining patterns were each set to be missing in 20% of the samples (N samples = 10). The proportion of missing values were set to 10%. We then used the `ampute` function to introduce MAR missing values according to the described scheme. An illustration of the weights used for selection of peptides for amputation is given in Figure S1.

For MNAR generation, we selected 9% peptides from mid-range abundance. We selected all other peptides from the same protein as the selected peptides. We partitioned the selections into four patterns and set them to be missing across samples at the frequency of 0.6,0.2,0.1 and 0.1, according to the MNAR mechanism. The proportion missing was set to 1%, hence each sample had 1% MNAR missing values. We selected peptides with mid-range abundance and all other peptides in the protein group of selected peptides to mimic dropout patterns in DDA, where peptides from the same protein are detected at high abundance in one experimental group, but are missing in the other experimental group due to LOQ. The patterns and their corresponding frequencies were deliberately determined such that peptides from the same protein are amputed in at least one experimental group, and remain intact in other experimental groups.

#### 2.1.3 Missing value imputation methods

We benchmarked `msImpute` against six established imputation techniques in proteomics, namely Zero replacement (*22*), MinDet (*23, 24*), Perseus-style imputation (*25*), QRILC (*26*), K-Nearest Neighbors (KNN) (*27*) and MLE (*28,29*). MinDet replaces missing values in each sample with a minimal observed value in that sample, determined by the *q*^*th*^ quantile of the observed values. Perseus replaces missing values by random numbers drawn from a normal distribution with a width of 0.3 and down shift of 1.8. We deployed Perseus in column-wise imputation mode (the default mode), where imputation is applied to each expression column separately. QRILC imputes missing values by random draws from a truncated normal distribution with parameters estimated using quantile regression. In KNN, missing values in a sample are replaced by the average of k closest observed measurements in that sample (local estimation), whereas in MLE missing values are replaced by Maximum Likelihood estimates (global estimation).

MinDet and Zero replacement are known as single value approaches, and as with QRILC and Perseus, are commonly used for imputation of left-censored MNAR data. KNN is known as a local similarity approach, and MLE is a global similarity approach to imputation. We used *imputeLCMD* (*26*) implementation of these imputation methods with default parameters. We re-implemented Perseus’s (column-wise) imputation C++ script in R, where missing values were replaced by random draws from a multivariate normal distribution, using the `rmvnorm` function in *mvtnorm* R package. The multivariate normal distribution was parameterized by the following (vector) of means ***µ*** and standard deviations ***σ***:

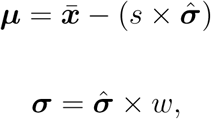

where 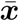 denotes sample means, 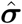 denotes sample standard deviations, *s* and *w* denote shift and width parameters, respectively. We set *s* = 1.8 and *w* = 0.3, as per defaults of Perseus.

### 2.2 Performance evaluation

We used Root Mean Squared Error (RMSE) to quantify the error introduced in intensity values by imputation in simulated datasets. Let (*i, j*) denote peptide *i* in sample *j, x*_(*i,j*)_ the ground truth log_2_ intensity and denote 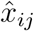 the corresponding imputation. Let 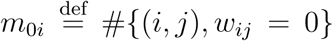 denote total number of missing values for *i*th peptide. The RMSE for the *i*th peptide is defined as:

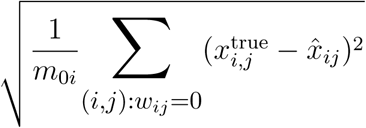

We looked at imputation error against average intensity of the peptides to investigate if imputation error varies as average intensity increases. We also evaluated the impact of imputation on results of differential expression analysis, and perturbation of global and local structures.

#### 2.2.1 Differential peptide expression

We used linear models with Empirical Bayes moderated t-statistics implemented in *limma* (*30*) R/Bioconductor package to identify peptides differentially expressed between different concentration of spiked-ins in benchmark datasets. Peptides were called differentially expressed if they achieved a False Discovery Rate of 0.05. We evaluated the performance of imputation methods by number of spiked-in peptides detected in top N DE peptides, number of True Positives (peptides from spiked proteins correctly called differentially expressed), False Positives (peptides falsely called DE), Precision (number of peptide from spiked proteins correctly called DE from all discoveries). In addition, for each method we evaluated the rank of truly differentially expressed peptide in the top most significant findings against the false discoveries made by the method.

#### 2.2.2 Perturbation to global and local structures

We quantified distortion to local and global structures post imputation. We define local structure as distance between biological (or technical) replicates of an experimental condition, and global structure as distance between biological groups in an experiment. We use four metrics to quantify such distortions: KNN, KNC, CPD and Gromov-Wasserstein distance.

- **KNN** The fraction of *k*-nearest neighbor runs in the original data that are preserved as *k*-nearest neighbors in imputed data (*31*). KNN quantifies preservation of the local, or microscopic structure.
- **KNC** The fraction of *k*-nearest class means in the original data that are preserved as *k*-nearest class means in imputed data (*32*). KNC quantifies preservation of the mesoscopic structure after imputation.
- **CPD** Spearman correlation between pairwise distances in the original data and imputed data (*33*). CPD quantifies preservation of the global structure.
- **GW** Gromov-Wasserstein distance is a distributional divergence metric (*34*) that quantifies dissimilarity of local and global structures simultaneously. We compute the GW distance between Principal Components (PCs) of the imputed data and original data, where PCs in the original data are computed using peptides with high biological variance.

In addition to simulation studies, we assessed RMSE and the three metrics of structural properties (KNN, KNC and CPD) empirically in datasets where alternative measurements were available for missing peptides. We identified a matched DIA-TMT dataset (PXD011691), and a matched DDA-DIA (PASS00589). The latter is an example of MNAR as the intensity values tend to be missing because of LOQ in DDA, while the earlier is an example of MAR/MCAR. For each pair of datasets, we treated the dataset with more complete measurements as the ground truth. For the DDA-DIA pair we evaluated imputation on peptides missing in DDA that were detected in DIA, and for the DIA-TMT pair we carried out the assessments on peptides missing in DIA that were measured in TMT dataset. The datasets where each centered to ensure the magnitude of the error is comparable. We used *k*=3 nearest neighbor runs/classes for simulated datasets. For empirical evaluations we used *k*=3 for DIA-TMT and *k*=2 for DDA-DIA. The GW distance was computed for all datasets used in differential expression analyses using all PCs in imputed and original data. The PCs in original data were computed on top 1000 highly variable peptides in PXD011691, and top 50 in all other datasets. The highly variable peptides were determined by decomposition of the total variance into biological and technical variance similar to the *scran* package (*35*).

#### 2.2.3 Linearity of observed UPS2 protein abundances after imputation

The UPS2 consists of 48 proteins organized in 6 tiers of abundance. We assessed if imputation would preserve the linearity of observed protein abundances with the theoretical concentration of the spiked UPS2 proteins in a Top3 analysis. The average intensity of the three (or fewer, if fewer peptides were detected) peptides with the highest intensity, Top3, was determined for each protein detected (*36*). Peptides were chosen separately for each sample, so the same peptides were not necessarily used across the 10 samples. We used UPS2 FLI-DIA data in PXD011691 for the Top3 analysis. There was a total of 12 UPS2 proteins detected in this study. Our evaluations are based on 10 of 12 UPS2 proteins that passed our filtering steps and were retained in DE analysis of this dataset.

## 3 Results

### 3.1 Imputation by low-rank approximation via Alternating Least Squares

Any high-dimensional dataset *X*_*m×n*_ with *m* features and *n* observations can be approximated and reconstructed by a number *r* ≤ min(*m, n*) of linear combination of its features. This is the basis of the Singular Value Decomposition (SVD) technique widely used for pattern extraction. Founded on `softImpute-ALS` algorithm (*37*), `msImpute` finds a low-rank approximation of the incomplete peptide abundance matrix and reconstructs the peptide intensity matrix as the product of two low-rank matrices.

Let *X*_*m×n*_ denote the peptide intensity matrix with missing values where *m* denotes the number of peptides and *n* denotes the number of samples. Denote the indices of non-missing observations by the set Ω. `softImpute-ALS` combines Nuclear-Norm-regularized matrix approximation and maximum-margin matrix factorization to find two low-rank *r* ≤ min(*m, n*) matrices *A*_*m×r*_ and *B*_*n×r*_, such that the incomplete matrix can be reconstructed by the product of the two matrices, i.e. *X* ≈ *AB*^*T*^. The two matrices *A* and *B* are found by minimizing the following objective function:

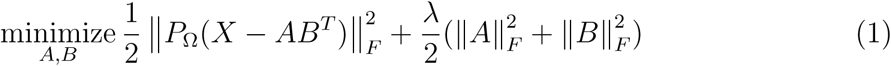

where *P*_Ω_ is the subset of observed peptide intensities, 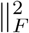 is the nuclear norm that encourages low-rank solutions, and *λ* is a shrinkage operator that controls the rank of the matrices being estimated. That is, we find two matrices A and B of lower dimensions (rank) than the measured peptide intensities, X, such that their products approximate X over the observed values with a reasonable accuracy (hence, the difference between X and 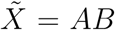 becomes negligible, for observed entries of X). The solutions are found by alternating between two Least Squares problems given in equations (2) and (3).

The matrix *A* = *UD* is initialized by random matrix *U*_*m×r*_ with orthonormal columns and *D* = *I*_*r*_, the identity *r × r* matrix. Given A, solve for B:

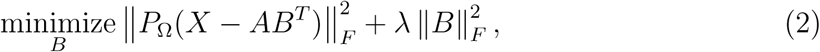

This is a multiresponse ridge regression with solution:

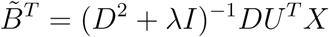

*B* = *V D* is reconstructed from SVD of 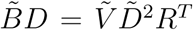, where 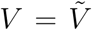 and 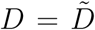. Given B, A is solved by

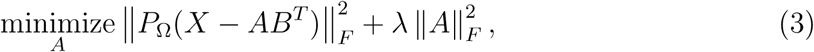

which is also a multiresponse ridge regression with solution

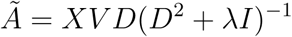

*A* is then updated by the product of two matrices *A* = *UV*, where *U* = *Ũ* and 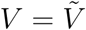 are estimated from SVD of 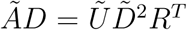. These steps are repeated until the difference between successive estimates of *AB*^*T*^ becomes negligible (i.e. the algorithm is converged). The parameter *λ* controls the rank *r* of *A* and *B* matrices, hence ensures the solution to equation (1) is low-rank. As *λ* decreases, the rank of solutions tend to increase.

`msImpute` first standardizes rows and columns of the matrix with missing values to have zero means and unit variances. Since missing values are present, mean and standard deviations cannot be estimated directly. `msImpute` first scales the data using the `biscale` algorithm proposed in (*37*), which estimates mean and variance of rows and columns using methods of moment. `msImpute` first scales the data using this algorithm. It also computes the optimal, data-driven value for *λ* from the scaling step, then finds the low-rank approximation 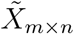 of *X*_*m×n*_ by estimating *Ã* and 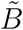. The complete peptide intensity matrix is then reconstructed by 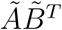. We recommend users apply `msImpute` to filtered, normalized, log_2_-transformed, peptide intensity values.

### 3.2 Assessment of imputation error

Root Mean Squared Error is commonly used as a measure of imputation error in proteomics benchmark studies (*12, 13, 22*). One expects the error to be uniform for peptides at all ranges of abundance. Hence, absence of any trends in RMSE versus average peptide intensity is desired, if an appropriate imputation procedure is adopted.

Evaluation of RMSE as a function of mean intensity in simulated MAR (Figure 1a), and NMAR data (Figure 1b) suggested that in MAR datasets the error is smaller and relatively uniform over the average abundance range in MLE and `msImpute`, whereas KNN tends to make large errors for both low- and high-abundance peptides. Although the methods developed for left-censored MNAR missing data, MinDet, QRILC, Perseus, and Zero, tend to maintain a uniform small (with the exception of Zero) error in MNAR data, they result in largest error when applied to MAR data. The imputation error is monotone increasing over average log_2_ intensity for Zero replacement. It has an unbalanced V-shape trend for MinDet, QRILC and Perseus, where imputation error is large for both low and high abundance peptides, but is larger for peptides at high abundance. Replacement of missing values by zero resulted in largest imputation error in both MAR and MNAR data. Similar trends were identified in our empirical evaluations of RMSE and mean intensity in MAR and MNAR datasets where alternative measurements were available for missing values (Figure S2a).

**Figure 1:**
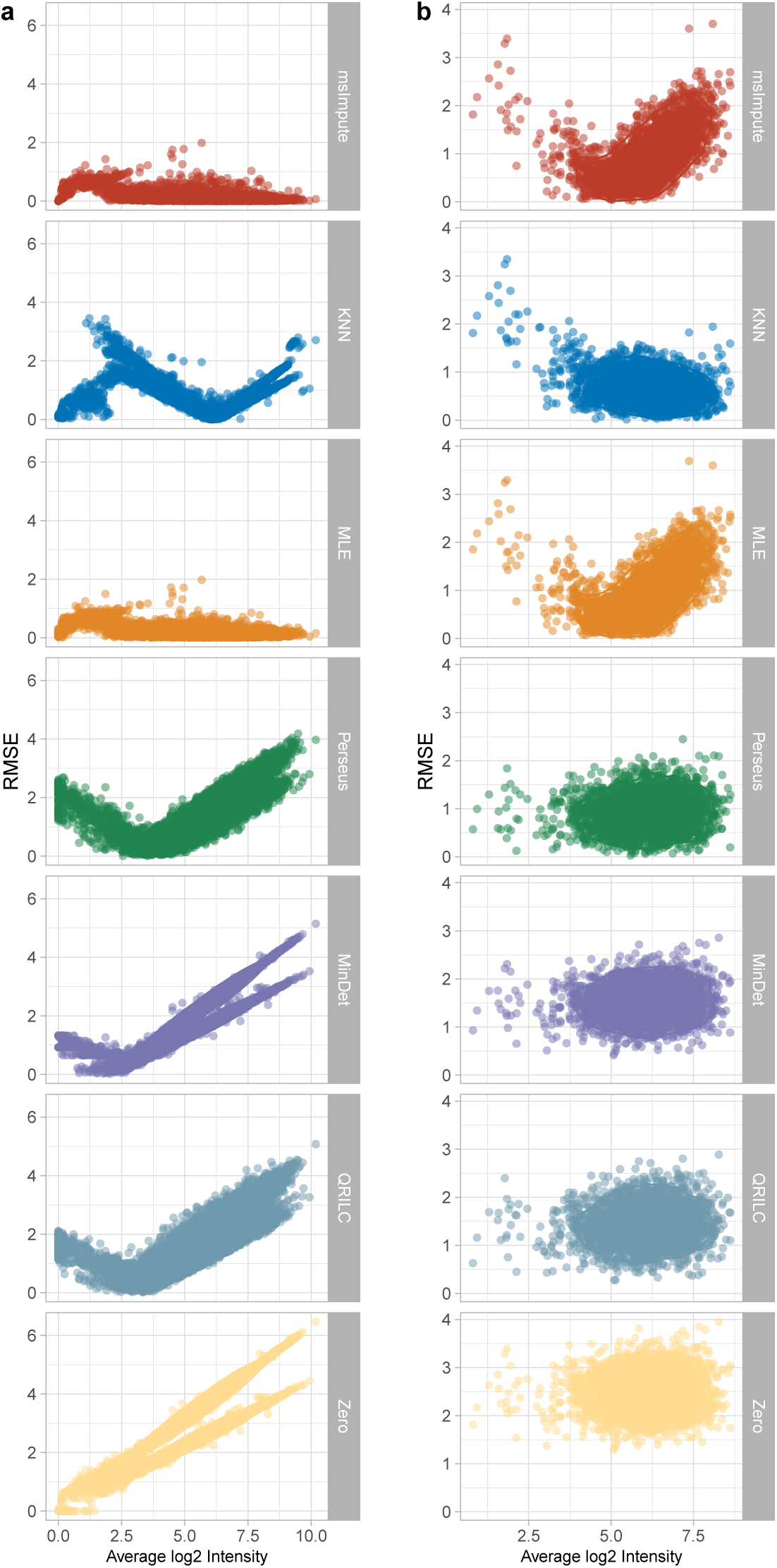
Evaluation of Imputation Error. RMSE versus Average intensity in simulated MAR dataset **(a)**, and MNAR dataset **(b)** by imputation procedure. Small errors and uniform trends in RMSE versus mean intensity are desired.

### 3.3 Assessment of the impact of imputation on Differential Expression

We found that `msImpute` tends to behave similar to MLE and KNN in recovering true differentially expressed peptides and outperforms single value or left-censored MNAR imputation approaches such as replacement with zero, MinDet, QRILC and Perseus in PXD011691 and PXD002370 datasets (Figure 2). `msImpute` detects a larger number of UPS1/2 peptides in top differential expression calls (Figure 2a), orders spiked-in peptides at the top DE discoveries while making less false discoveries (Figure 2b), and maintains highest precision and low false discovery rate (Figure 2c,d) compared to single value and left-censored MNAR approaches in these datasets. We observed a favorable performance when no imputation was applied to datasets in Figure 2 that is similar to imputation by MLE. This was, however, coupled by hundreds of peptides being discarded from the computations, as their fold-change or variances could not be estimated by *limma*. In addition, we observed that the UPS2 protein abundances, measured as the average of top 3 most intense peptides (i.e. Top3) across replicates, in all imputation methods except Zero replacement maintain a linear association with theoretical concentrations in PXD011691 dataset (Figure S3).

**Figure 2:**
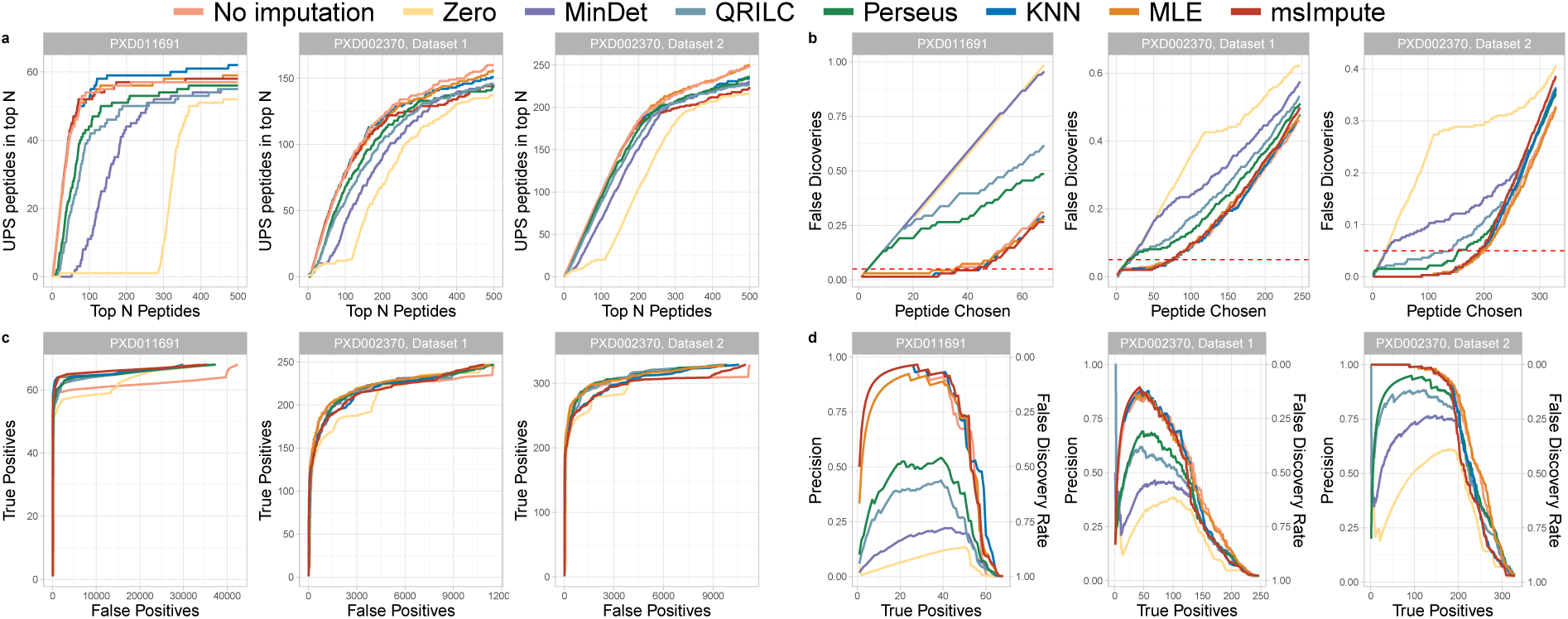
Impact of imputation on Differential Expression Analysis at the peptide level in MAR data. **(a)** Number of peptides from UPS/spiked-in proteins in top N differentially expressed peptides per imputation method in six benchmark datasets. A large number of UPS peptides detected in top N discoveries is desired. **(b)** Ranking of the UPS/spiked-in peptides in top N differential expression calls and false discoveries made by each imputation technique in benchmark datasets. An ideal imputation procedure is expected to order UPS/spiked-in peptides at the top N discoveries, while maintaining small number of false discoveries.**(c)** True Positives (UPS/spiked-in peptides correctly called DE) vs False Positives (none UPS/spiked-in peptides mis-identified as DE) in benchmark datasets with MAR-type missing values. **(d)** Precision versus True Positives. Proportion of True Positives (peptides from UPS/spiked-in proteins correctly detected as differentially expressed) from all differential expression calls versus number of True Positives per imputation procedure in six benchmark datasets. The secondary axis captures False Discovery Rate (FDR). High precision and small FDR are desired in DE analysis.

We also observed that `msImpute` can result in high false discovery rate and fail to recover a considerable number of true DE peptides (i.e. true positives) in some datasets (Figure 3a-d), where the evaluations were in favor of left-censored MNAR approaches, particularly QRILC and Perseus. We reasoned that the variations in the performance of `msImpute` in differential expression analyses is linked to the nature of missing values in datasets.

**Figure 3:**
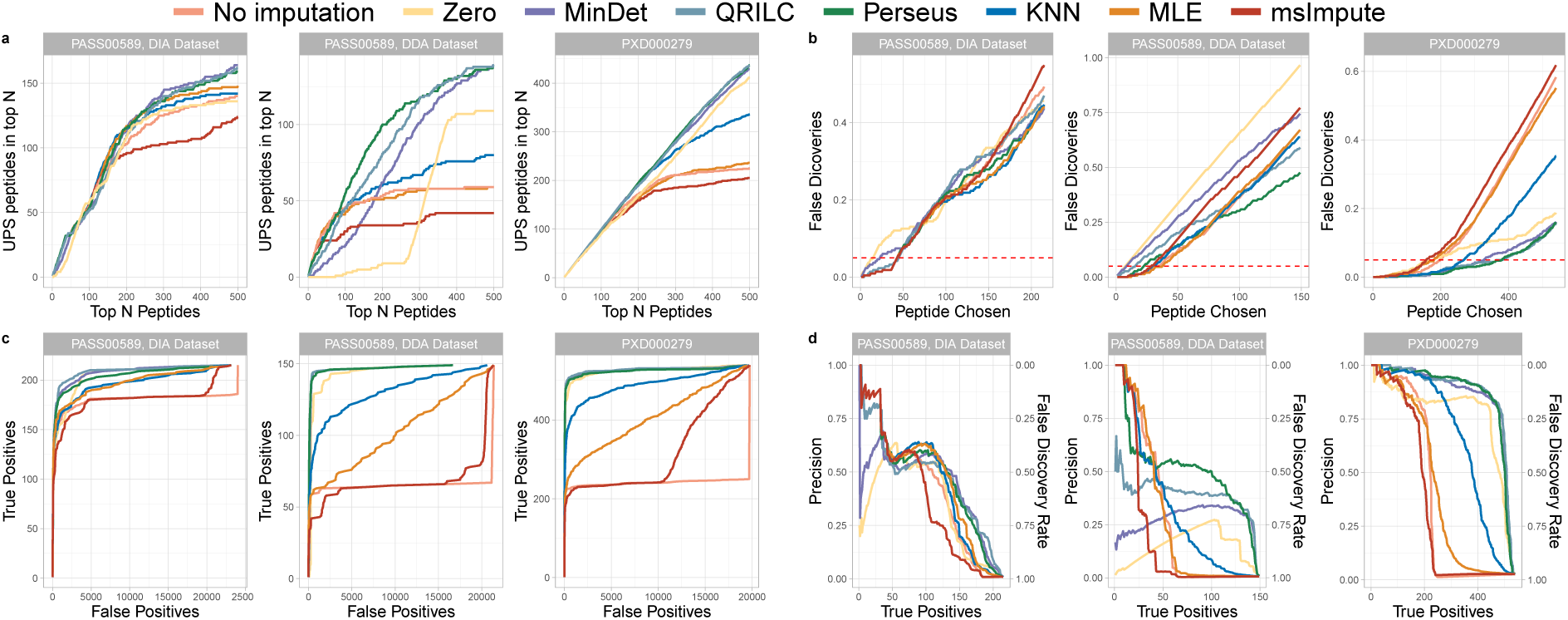
Impact of imputation on Differential Expression Analysis at the peptide level in MNAR data. **(a)** Number of peptides from UPS/spiked-in proteins in top N differentially expressed peptides **(b)** Ranking of the UPS/spiked-in peptides in top N differential expression calls and false discoveries made by each imputation technique **(c)** True Positives (UPS/spiked-in peptides correctly called DE) vs False Positives (none UPS/spiked-in peptides mis-identified as DE) **(d)** Precision versus True Positives.

### 3.4 A visual diagnosis for MAR/MNAR

We developed a visual test for MAR/MCAR and MNAR diagnosis based on dropout patterns of peptides with high biological dropout rate. We fit a linear model to peptide dropout rate, *D*, against peptide average intensity, *µ*:

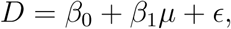

where *ϵ* is the error term. The *k* peptides with largest residual have high dropout rate and are highly expressed. If such peptides tend to be missing completely in one biological group, then the dropout probability of a missing peptide depends on other missing peptides, which is, by definition, describing the MNAR mechanism. We refer to top *k* peptides with largest residual as *High Biological Dropouts (HBD)*.

We identified top 500 high biological dropout peptides as described earlier in each benchmark dataset and produced heatmaps of their detection pattern (Figure 4). In datasets where `msImpute` failed to maintain reasonable false discovery and true positive rate, we noticed the presence of large blocks where intensity values were measured completely in one experimental group, but were missing in the other group, for example, due to LOQ. This indicates that the dropout probability of a missing peptide depends on other missing peptides in closely related samples. By definition, such missing values are MNAR, hence why left-censored MNAR imputation methods outperform in this data. We, therefore, inferred that data in Figure 4b are MNAR, whereas data in Figure 4a are MAR/MCAR as there is no systematic pattern in missingness patterns of high biological dropout peptides. We suggest users produce heatmaps of dropout patterns of *HBD* peptides as in Figure 4. If blocks of missing peptides are observed in one biological group, the underlying missingness mechanism is MNAR. Otherwise intensity values are MAR in which case `msImpute` will outperform imputation techniques that replace missing values with a determined small value, or random draws from a Gaussian distribution. Peptides with high biological dropouts can be identified using `selectFeatures` function in *msImpute* software.

**Figure 4:**
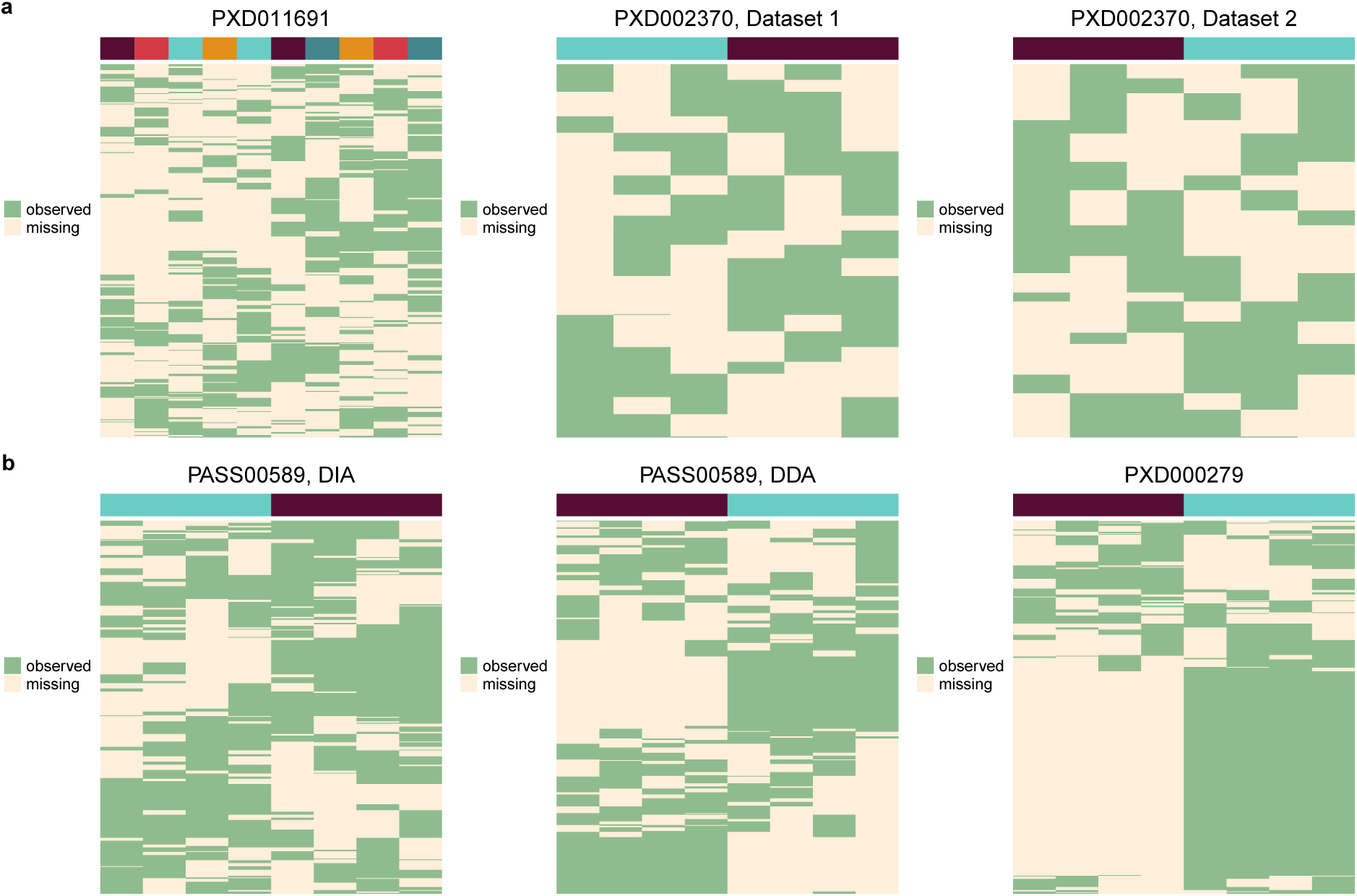
Performance of an imputation procedure depends on the nature of missingness in the dataset. Heatmap of detection patterns (1 if observed, 0 if missing) of high biological dropout (HBD) peptides in MAR **(a)** and MNAR datasets **(b)**. In MNAR data, dropout probability of a peptide depends on other missing peptides and dropout often occurs in a group of similar samples. Hence, MNAR can be identified by detection of block of missing values (i.e. related peptides from the same protein missing in one biological/experimental group). However, the block-wise missing structures are absent in MAR settings. Rows are peptides and columns are samples. Columns are annotated by experimental group.

### 3.5 Preservation of global and local structures post-imputation

Using both simulated and real MAR/MNAR data, we could demonstrate that imputation introduces distortions to local (within experimental group) and global (between experimental groups) similarities in a dataset. While perturbation to local structures is related to variance of measurements, distortion in global structures pertains to accuracy of fold-changes estimates between experimental groups. We observed that `msImpute` and KNN achieve lowest average RMSE in simulated MAR (Figure 5a) and NMAR (Figure 5b) data, respectively. Although low average RMSE indicates small deviation of imputed values from ground truth, it does not guarantee preservation of pairwise distances between samples, within or between experimental conditions.

**Figure 5:**
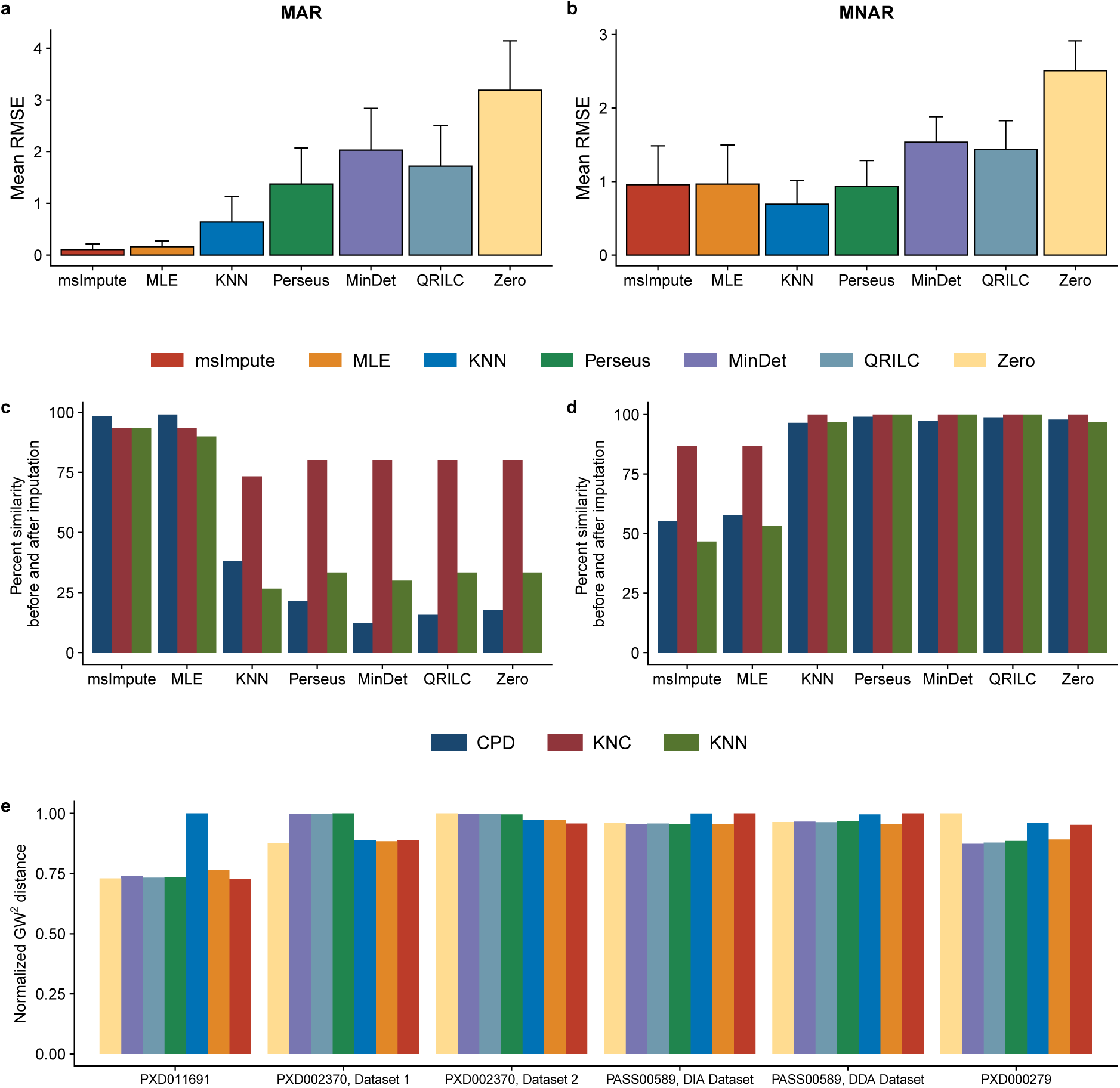
Mean RMSE and distortion to local and global structures in simulated data. Average RMSE (averaged over all missing peptides) in MAR **(a)** and MNAR **(b)** setting. Percentage of preserved k-nearest neighbor runs (samples) and class means, and the pair-wise Pearson correlation between runs in original and imputed data in MAR **(c)** and MNAR **(d)**. Squared Gromov-Wasserstein distance for benchmark datasets used in differential expression analyses. For each dataset we normalize the distances by the maximum *GW* ^2^.

While all imputation methods preserve mesoscopic structures (similarity of related samples) reasonably well in both MAR (Figure 5c) and MNAR (Figure 5d) datasets, preservation of local and global properties by various imputation methods depend on the underlying missing data generation mechanisms. In MAR data, single value approaches MinDet, Zero and left-censored MNAR methods only maintain, on average, 30% of global and local structures, while these structures are almost perfectly preserved by `msImpute` and MLE. Local similarity approaches to imputation can preserve 40% of global and local structures in MAR data (Figure 5c). In MNAR data, the fraction of preserved nearest neighbors and correlation between pairwise distances are lower in data imputed by MLE and `msImpute`, whereas an almost perfect preservation of local and global similarities is observed by KNN, Zero, MinDet, QRILC and Perseus (Figure 5d). Our empirical evaluation of preservation of the structures in the MAR and NMAR data confirms similar conclusions to simulated datasets (Figure 5e and Figure S2b,c).

In the six datasets that we used to benchmark performance in differential expression analyses, we computed the Gromov-Wasserstein distance between imputed data and the original data. This distance metric measures preservation of local and global structures simultaneously, so that larger distortions to the data post imputation result in larger distances between imputed and original data. We observed that in MAR datasets global similarity approaches, and in MNAR datasets left-censored missing approaches tend to result in smaller distance estimates, hence better preservation of local and global structures (Figure 5e).

## 4 Discussion

Many proteomics data analysis approaches such as differential expression, Network Analysis, classification and multi-omics integration require complete observations and non-NA measurements. Missing values are inherent in proteomics datasets. Hence, imputation of missing values is an essential part of data processing.

We evaluated the performance of the proposed imputation method along with the widely used local, global and left-censored NMAR missing imputation approaches. Evaluation of RMSE of imputed peptide log-intensities as a function of mean log-intensity in simulated MAR and NMAR data suggested that MLE and `msImpute` have similar imputation error. Similarly, left-censored MNAR methods produce similar RMSE trends. RMSE is on average lower in `msImpute` compared to all other methods in simulated MAR data, and is smaller than the left-censored missing, single value imputation, or Zero replacement approaches in simulated NMAR settings. We observed that, regardless of the nature of missing data, imputation by zero results in largest error/deviation from ground truth. We note that `msImpute` is akin to global similarity approaches of imputation such as MLE, and outperforms single value and left-censored MNAR approaches, that are mostly based on random sampling from a normal distribution, in MAR data.

In differential expression analyses, `msImpute` can reliably detect and rank true DE peptides at the top table of the findings in MAR data. The benchmark differential expression results are in favor of QRILC and Perseus in MNAR data. Under MAR settings, `msImpute` and MLE preserve data structures post imputation. Replacement of missing values by zero, or sampling from a Gaussian distribution are common choices of imputation by the proteomics community and here we demonstrate that mean RMSE and perturbations to local and global structures are generally very large for these methods in MAR data. Hence, these imputation strategies should be practiced with caution. Our benchmark and simulation studies suggest that under a MAR assumption `msImpute` and MLE, and under MNAR assumption QRILC and Perseus are safe choices for imputation. Importantly, missing values can be left untreated in differential expression analyses if they occur randomly, and indeed we showed that performance is comparable to that of imputation by MLE. However, many peptides will be discarded when linear models are fitted, as their fold-change or variance cannot be estimated, which results in the loss of information from the data.

Our evaluations suggest that there is a resemblance in performance among methods in each category of imputation procedures (e.g., local-similarity, global-similarity, left-censored MNAR). Therefore, based on the datasets used in this study, there is likely to be one dominant pattern of missingness in datasets. Our proposed method for MAR/MNAR pattern identification based on peptides with high biological dropout rate can, hence, serve as an effective visual tool to infer the governing pattern. For each sample in a dataset, the missingness patterns of *High Biological Dropout peptides* introduced in this work can be encoded as a binary vector, where 1 denotes the measurement for the peptide is missing in the sample and 0 denotes otherwise. Such encoding results in a binary matrix of dimensions *n × k*, where n is the number of samples and *k < m* is the number of top *HBD* peptides, that can be used to train a supervised classification models such as Logistic Regression or Support Vector Machine (SVM) to predict the biological group of the samples. A Receiver Operator Characteristic (ROC) value larger than 0.5 implies that the missing patterns contain information that is predictive of biological groups, and hence, a Missing Not At Random mechanism. `msImpute` has been successfully applied in previous proteomics studies (*38*).

## Abbreviations

DE: Differential Expression
MAR: Missing At Random
MCAR: Missing Completely At Random
MNAR: Missing Not At Random
SVD: Singular Value Decomposition
FDR: False Discovery Rate
MS: Mass Spectrometry
LC-MS: Liquid Chromatography - Mass Spectrometry
DDA: Data-dependent Acquisition
DDA: Data-independent Acquisition
ALS: Alternating Least Squares
RMSE: Root Mean Squared Error
HBD: High Biological Dropout
LOQ: limits of quantification
PC: Principal Components

## Acknowledgments

The authors would like to thank Professor Gordon Smyth from WEHI for his comments on the proposed imputation method, and advice on adapting *limma* for differential expression analysis in proteomics data. We also thank Giuseppe Infusini, Laura Dagley, Jarrod Sandow, and Samantha Emery from WEHI, Anup Shah (Monash University) and Jeremy Potriquet (SCIEX Brisbane) for helpful discussions.

## Data and software Availability

Peptide-level quantification (in tab-delimited format) for the benchmark datasets were obtained from ProteomXchange using accession numbers PXD011691, PXD002370, PASS00589 and PXD000279.

The *msImpute* software is a R package available for download from https://github.com/DavisLaboratory/msImpute [DOI: 10.5281/zenodo.3980269]. The KNN, KNC, CPD and GW metrics used in this manuscript for the assessment of post imputation preservation of structures are also included in the package.

## Supplementary materials

**Figure S1:**
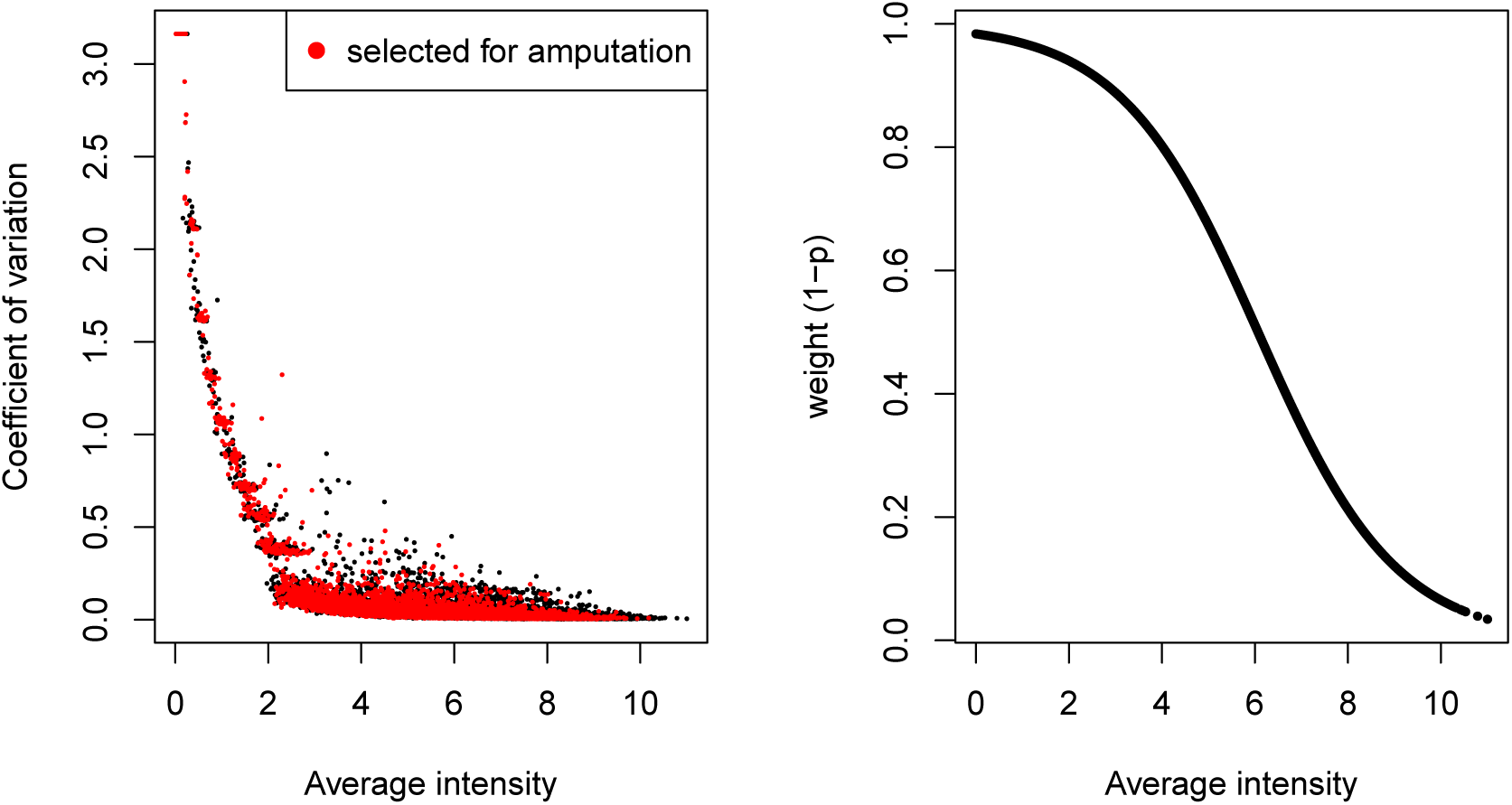
Peptides are selected for amputation according to a weighted sampling procedure. The sampling procedure employed reflects the mean-dropout trend commonly observed in proteomics data, where low abundance peptides have higher dropout rate. Dropout rate decreases for peptides with high average intensity Right Each peptide is assigned a weight according to its average log_2_ intensity during sampling. Left Coefficient of variation versus average log_2_ intensity for all peptides and peptides selected for amputation according to the weighted sampling scheme.

**Figure S2:**
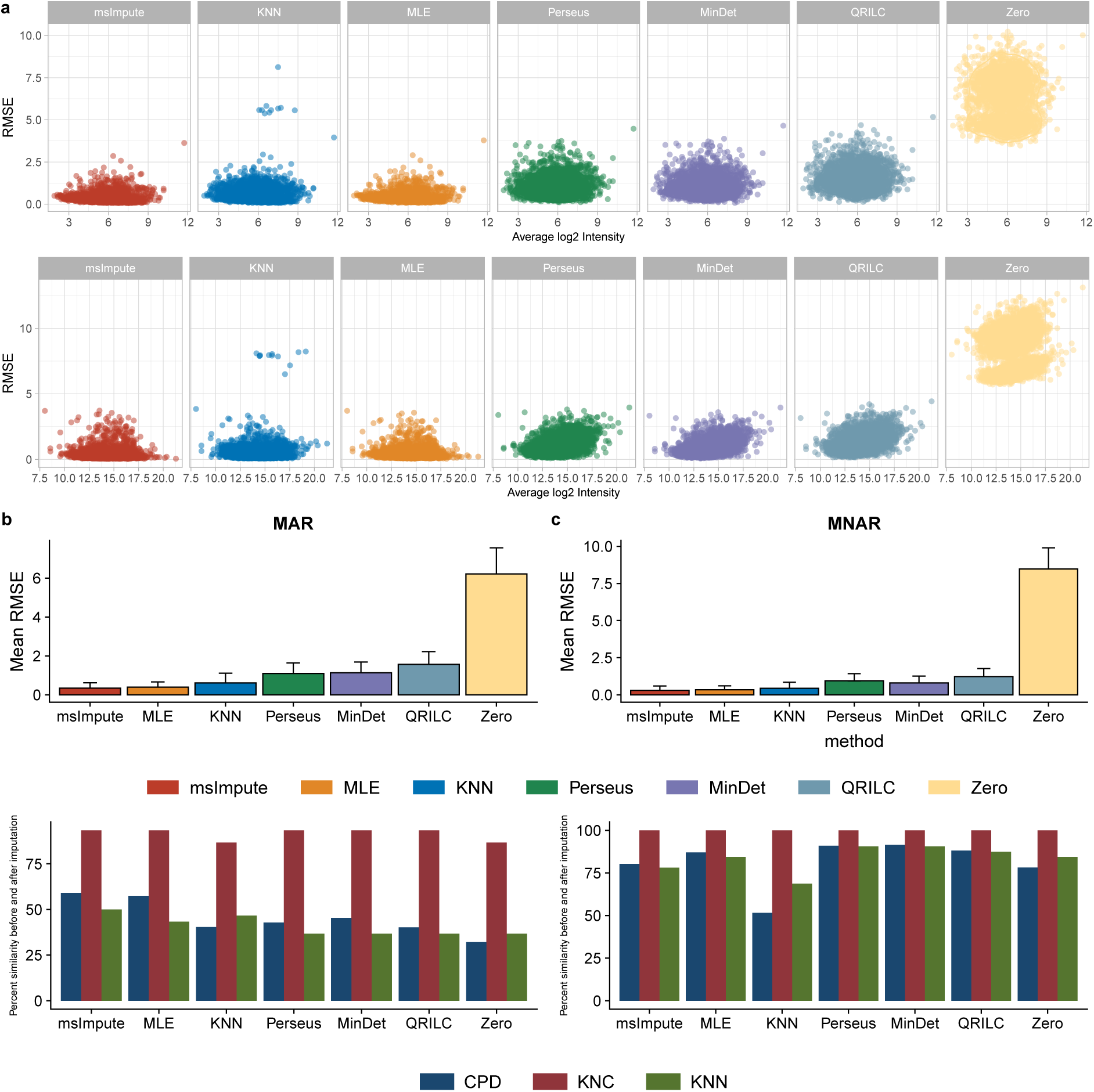
Empirical evaluation of RMSE and distortion to local and global structures in real data. RMSE versus Average intensity in simulated MAR (PXD011691 DIA) dataset **(a, upper panel)**, and MNAR (PASS00589 DDA) dataset **(a, lower panel)** by imputation method. Average RMSE (averaged over all missing peptides) in MAR **(b, upper panel)** and MNAR **(c, upper panel)** setting. Percentage of preserved k-nearest neighbor runs (samples) and class means, and the pair-wise Pearson correlation between runs in original and imputed data in MAR **(b, lower panel)** and MNAR **(c, lower panel)**.

**Figure S3:**
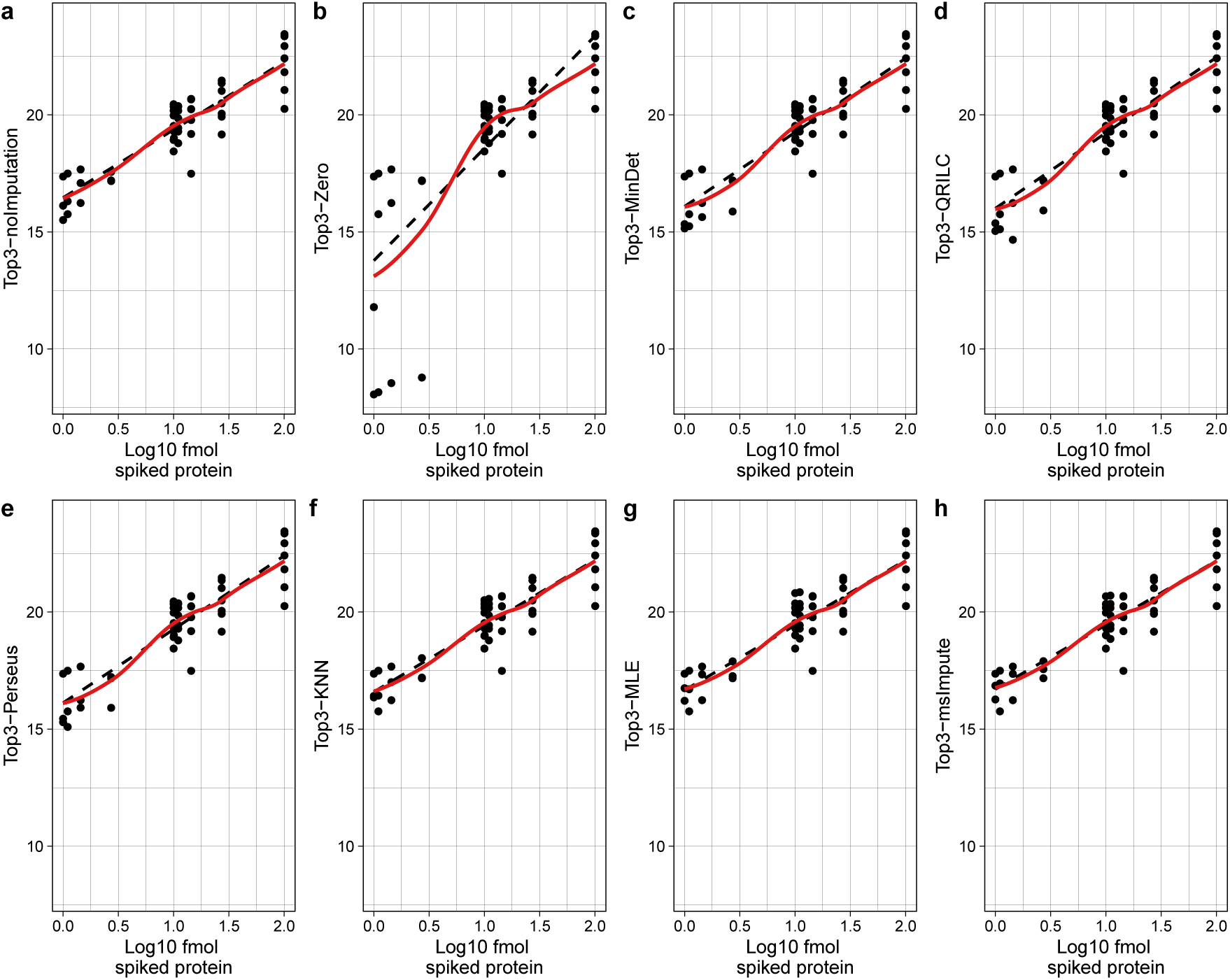
Top3 analysis. Linearity of observed UPS2 protein abundances with theoretical concentrations is maintained by all imputation approaches, except Zero replacement. Black dash line is the linear fit to the data points, red solid line is a loess (non-linear) fit. For all methods except Zero replacement, the linear and non-linear fits overlap, which indicates that overall the observed UPS2 protein abundances maintain linearity with theoretical concentrations.

## Notes

### Competing Interest Statement

The authors have declared no competing interest.

